# ITRAQ-based quantitative proteomic analysis of *japonica* rice seedling during cold stress

**DOI:** 10.1101/2021.08.19.457014

**Authors:** Dongjin Qing, Yinghua Pan, Gaoxing Dai, Lijun Gao, Haifu Liang, Weiyong Zhou, Weiwei Chen, Jingcheng Li, Juan Huang, Ju Gao, Chunju Lu, Hao Wu, Kaiqiang Liu, Guofu Deng

**Author notes:** Correspondence (K.L.); (G.D.); (K.L.); (G.D.).

## Abstract

Low temperature is one of the important environmental factors that affect rice growth and yield. To better understand the *japonica* rice responses to cold stress, isobaric tags for relative and absolute quantification (iTRAQ) labeling based quantitative proteomics approach was used to detected changes in protein level. Two-week-old seedlings of the cold tolerance rice variety Kongyu131 were treated at 8°C for 24, 48 and 72 h, then the total proteins were extracted from tissues and used for quantitative proteomics analysis. A total of 5082 proteins were detected for quantitative analysis, of which 289 proteins were significantly regulated, consisting of 169 uniquely up-regulated proteins and 125 uniquely down-regulated proteins in cold stress groups relative to control group. Functional analysis revealed most of regulation proteins involved in photosynthesis, metabolic pathway, biosynthesis of secondary metabolites and carbon metabolism. Western blot analysis showed that protein regulation was consistent with the iTRAQ data. The corresponding genes of 25 regulation proteins were used for quantitative real time PCR analysis, and the results showed that the mRNA level was not always parallel to the corresponding protein level. The importance of our study is providing new insights into cold stress responses in rice on proteomic aspect.

## Introduction

Rice (*Oryza sativa*) is one of the most important food crops in the world, feeding about half of the population [1]. Climate condition is an important factor affecting rice yield, especially low temperatures as a common environment factor affect the whole life cycle of rice growth in tropical and subtropical areas, ranging from vegetative to reproductive stages [2]. Low temperature can cause seedlings of cold-sensitive rice cultivars severe injury in the early season, and reduce growth rate, pollen sterility in the late season [3, 4]. Low temperature stress occurs frequently and has wide influence range in the word with the increasing of global climate anomalies [5, 6]. Therefore, it is important to explore more low temperature regulation genes/proteins for understanding the cold-tolerance mechanism and breeding cold-tolerant rice cultivars.

An increasingly number of molecular genetic studies have elucidated how rice plants respond to low temperature stress and genes/proteins involved in the response. Low temperatures signal is first perceived by the temperature sensor COLD1/RGA1 complex on plasma membrane, then the complex triggers an influx of calcium, reactive oxygen species (ROS) production, ABA accumulation and MAPK cascade (OsMKK6-OsMPK3) reactions to active downstream transcription factors responses in the nucleus [7-9]. Recently, the vitamin E-vitamin K1 sub-network of COLD1 downstream pathway was found to be responsible for chilling tolerance divergence [10]. Other components of cold tolerance also have been identified recently, such as *CTB4a* interacts with a beta subunit of ATP synthase AtpB for mediating ATP supply in rice plant cells to improve cold tolerance[11], standing variation of cold tolerance gene *CTB2* and *de novo* mutation of *CTB4a* facilitate cold adaptation of rice cultivation from high altitude to high latitude areas [12]; bZIP73^Jap^ in *japonica* rice cultivars interactions with bZIP71 to modulates abscisic acid (ABA) levels and reactive oxygen species (ROS) homeostasis for enhancing rice tolerance to cold climate [13]; OsMADS57 interacts with OsTB1 and both directly target *OsWRKY94* and *D14* for rice adaptation to cold [14]. At rice seedling stage, the cold tolerance associated gene *qCTS-9* was found in hybrid rice under different cold environments using QTL mapping and genome-wide expression profiling methods [15]. *qPSST6*, found from cold tolerance *japonica* rice variety Kongyu131with QTL mapping and Seq-BSA approach, was validated to be a functional gene relative to cold resistance [16]. Three genes (LOC_Os01g55350, LOC_Os01g55510 and LOC_Os01g55560) [17] and 67 QTLs [18], which screened out by genome-wide association analysis (GWAS) were associated with cold tolerance of *indica* and *japonica* rice, respectively. In a RNA-seq comparative analysis of cold-stressed post-meiotic anther from cold-tolerant and cold-susceptible rice cultivars, a number of ethylene-related transcription factors were found to be putative regulators for cold responses [19].

Proteomic approach is a robust strategy for the large-scale identification of proteins, and it has been used for profiling proteins in rice recently [20]. Two-dimensional gel electrophoresis (2-DE) was used to separate proteins of rice with cold treatment, and cold response proteins were identified using mass spectrometry analysis in early proteomics studies [21-25]. iTRAQ is a powerful mass spectrometry technology, which can quantify proteins’ relative expression abundance by measuring relative peak areas of MS/MS mass spectra of iTRAQ-labeled peptides [26]. More and more cold response proteins in rice were monitored and characterized by iTRAQ-labeling approach. For example, differentially expressed proteins in cold stress treated rice involved in photosynthesis, metabolism, transport, ATP synthesis, ROS, stress response, DNA binding and transcription, cell growth and integrity, unknown function proteins were found using iTRAQ labeling coupled with LC-MS/MS [27-29]. However, some different cold response proteins were found in different rice cultivars by proteomic approach, and up to now, only a small number of cold-response proteins have been identified.

To better understand the cold tolerance mechanism of *japonica* rice, we employed iTRAQ labeling proteomics method to investigate the proteomic response of cold stress of *japonica* cold-resistant rice cultivar Kongyu131 in this study. Rice seedling tissues were harvested after exposed to 8 °C low temperature condition for 0, 24, 48 and 72 h. iTRAQ was used for quantifying relative protein abundance, and different expression proteins were obtained at each time point by comparing to the control samples. Our results show that a large portion of cold stress-regulated proteins were reported for the first time.

## Materials and methods

### Plant material and cold stress treatments

The japonica rice variety Kongyu131 which strongly resist to cold weather and widely planted in the northeast area of China was used in this study. Rice seedlings were grown in the growth chamber with a 16-h light (28°C)/8-h dark (25°C) condition for 2-week. Cold tress treatments were performed by decreasing temperature to 8°C, and collected tissues and frozen in liquid nitrogen at 0 h, 24 h, 48 h and 72 h respectively, then stored at -80°C refrigerator for protein extraction. For physiology experiment, 2-week-old rice seedlings were separated into groups and treated at 80°C for 4 days, then were transferred to the normal growth condition for 3 days for survival rate determination, the ratio of surviving plants to total plants was calculated.

### Protein extraction, digestion and iTRAQ labeling

Protein extraction was performed according to previous methods [54-56] with some modifications. The frozen rice seedling tissue (0.5 g) was ground to fine powder with -80 °C pre-cold mortar and pestle. The tissue powder was extracted with 5 volume (g/mL) of extraction buffer containing: 8 M urea,150 mM Tris-HCl, pH 7.6, 1.2% Triton X-100, 0.5% SDS, 5 mM ascorbic acid, 20 mM EDTA, 20 mM EGTA, 5 mM DTT, 50 mM NaF, 1 mM PMSF, 1% glycerol 2-phosphate, 1× protease inhibitor (complete EDTA free; Roche) and 2% polyvinylpolypyrrolidone. The extract was centrifuged at 110,000 ×g for 2 h at 10 °C to get rid of cell debris at the bottom of centrifuge tube. The total protein in supernatant was precipitated with 3 volumes of -20 °C pre-cooled acetone:methanol (12:1 v/v) for at least 2 hours. Collected the protein pellet by centrifugation at 11,000 ×g for 20 min and washed two times with acetone:methanol (12:1 v/v), then re-suspended the dry protein pellet in re-suspension buffer (100 mM Tris-HCl, pH 8.0, 8 M urea). The concentration of total protein was measured by Bradford method and the proteins were used for proteomics and western blot analysis.

The 200 μg protein samples were used for reduction reaction by adding 10 mM DTT and incubated at 56°C for 1 h, then followed by alkylation reaction by adding 40 mM iodoacetamide and incubated at room temperature for 30 min in dark condition. To digest protein with trypsin, urea was diluted below 2 M using 100 mM Tris-HCl (pH 8.0), then trypsin was added in the protein solution at 1:50 ratio (enzyme : protein, w/w) and incubated at 37 °C overnight. Acidized peptides by adding formic acid to end the digestion, then centrifuged at 12,000 ×g for 15 min, and the supernatant was subjected to peptide purification using Sep-Pak C18 desalting column. The peptide eluate was vacuum dried and stored at -20°C refrigerator.

For each sample, 100 μg peptide was used for iTRAQ labeling. The samples were separately labeled with different iTRAQ labeling reagents (113, 114, 115, 116) according to the manufacturer’s instructions. The labeled samples were mixed and subjected to Sep-Pak C18 desalting, then the complex mixture were fractionated using high pH reverse phase chromatography, and combined into 15 fractions. Each fraction was vacuum-dried and re-suspended in 0.1% formic acid for MS analysis.

### LC-MS/MS analysis, and protein quantification

LC-MS/MS detection was carried out on a hybrid quadrupole-TOF LC-MS/MS mass spectrometer (TripleTOF 5600, SCIEX) equipped with a nanospray source. Peptides were first loaded onto a C18 trap column (5 µm, 5 × 0.3 mm, Agilent Technologies) and then eluted into a C18 analytical column (75 μm × 150 mm, 3 μm particle size, 100 Å pore size, Eksigent). Mobile phase A (3% DMSO, 97% H_2_O, 0.1% formic acid) and mobile phase B (3% DMSO, 97% ACN, 0.1% formic acid) were used to establish a 100 min gradient, which comprised of: 0 min in 5% B, 65 min of 5-23% B, 20 min of 23-52% B, 1 min of 52-80% B, 80% B for 4 min, 0.1 min of 80–5% B, and a final step in 5% B for 9.9 min. A constant flow rate was set at 300 nL/min. For IDA mode analysis, each scan cycle consisted of one full-scan mass spectrum (with m/z ranging from 350 to 1500, ion accumulation time 250 ms) followed by 40 MS/MS events (m/z ranging from 100 to 1500, ion accumulation time 50 ms). The threshold for MS/MS acquisition activation was set to 120 cps for +2∼+5 precusors. Former target ion exclusion was set for 18 s.

Raw data from TripleTOF 5600 were analyzed with ProteinPilot (V4.5) using the Paragon database search algorithm and the integrated false discovery rate (FDR) analysis function. Spectra files were searched against the UniProt *japonica* rice reference proteome database using the following parameters: Sample Type, iTRAQ 8plex (Peptide labeled); Cys Alkylation, Iodoacetamide; Digestion, Trypsin; Quantitate, Bias correction, and Background correction was enabled for Specific Processing ; Search Effort was set to Rapid ID. Search results were filtered with unused score and false discovery rate threshold (FDR) at 1%. Decoy hits were removed, the remaining identifications were used for quantification. Proteins with a fold change of > 1.2 or < 0.83 and a p-value of < 0.05 were considered to be differentially expressed [57].

### Bioinformatics analysis of DEPs

All DEPs were used for hierarchical cluster analysis with Cluster 3.0 program. The DEPs were classified and grouped into different pathways according to Gene Ontology (GO) and Kyoto Encyclopedia of Genes and Genomes (KEGG). The protein-protein interaction networks were analyzed using the STRING 10 database (http://string.embl.de).

### Quantitative real time PCR and western blot analysis

To validate the MS quantification results at a transcript level, total rice seedling tissue mRNA extracted using TRIzol reagent (Invitrogen) was used for cDNA synthesis by using SuperScritRIII RT First Strand Synthesis Kit (Invitrogen) according to its protocol. SYBR® Premix Ex Taq ™ (Takara, China) was used for Real-time RT-PCR, and the specific primers (Table S2) for target genes amplification were designed using Primer Express 3.0 software. β-actin was used as an internal control gene. Three biological repeats were performed for each target gene in real-time RT-PCR.

Western blot analysis was performed according to Qing et al. with some modification [55]. The rabbit polyclonal antibody was raised against synthetic oligopeptides which were identified by MS, DAGDAAPPAAATTTER to make anti-A0A0N7KH91 polyclonal antibody. The peptide antibody was made commercially (GL Biochem Co., Ltd., Shanghai, China). The plant β-actin polyclonal antibody was bought from YIFEIXUE BIO TECH. The proteins used for western blot analysis were extracted from rice seedling tissue with urea extraction buffer, and separated on 15% SDS-PAGE gel and transferred onto a polyvinylidene fluoride membrane (Millipore, USA), which was probed with the anti-A0A0N7KH91 polyclonal antibody and anti-β-actin polyclonal antibody.

## Results

### Physiological response to cold stress

To validate contrasting stress phenotypes of Kongyu131 and other 11 cultivars in response to cold treatment, two-week-old rice seedlings were treated under 8°Cfor 4 days and recovered for 3 days. Before cold treatment, seedling plants of all varieties growth normally (Fig. 1A). After cold stress and recovery treatment, seedlings of *indica* rice varieties (Guanghui998, Jinweiai, 02428, Y58S and Dachangli) and *japonica* rice varieties (Liaoxing1, Liaoxing21 and Kunmingxiaobaigu) were completely wilted, whereas most seedlings of *japonica* rice varieties Kongyu131, Nipponbare and Daohuaxiang were able to survive (Fig. 1B). As seen in Fig. 1C, the survival rate of Kongyu131 after cold stress treatment was 77%, and is the highest survival rate by comparison to other varieties in this experiment.

**Fig 1.**
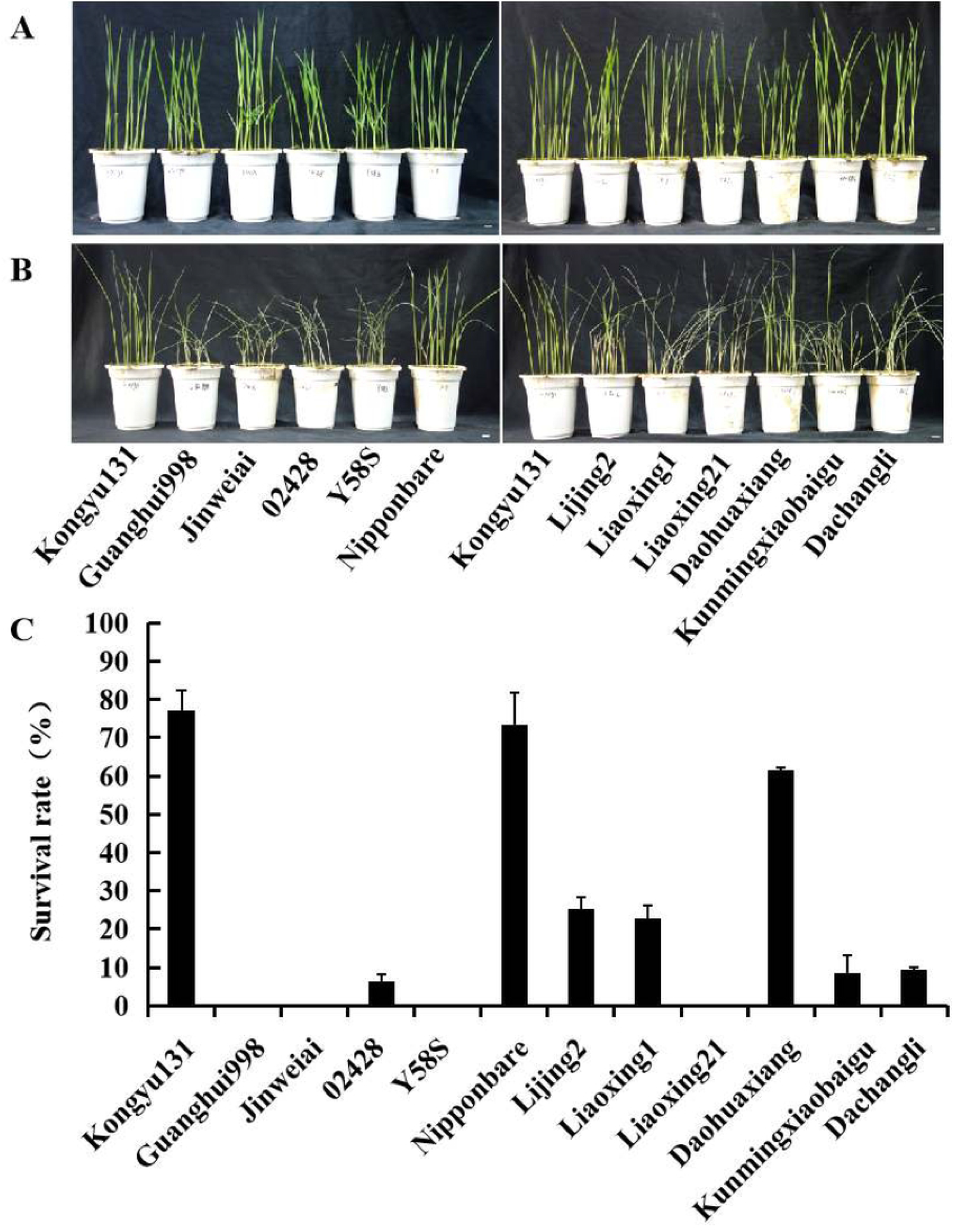
Phenotypes of low-temperature treated Kongyu131 and other rice cultivars after 3 days of recovery. Rice seedling were planted in a phytotron under light and temperature controls (the day/night cycle: 12 h with 28 °C and 12 h with 22 °C). 2-week-old rice seedlings were treated under 8 °C for 4 days and recovered for 3 days under normal growth conditions. A, Phenotypes of rice cultivars before low temperature treatment. B, Phenotypes of rice cultivars after 4 days of low temperature treatment and 3 days of recovery, bar = 1 cm. C, survival rate of rice cultivars after 3 days of recovery.

To understand which proteins have responses to cold stress in the Kongyu131 at the proteomic level, two-week-old seedlings planted in soil were subjected to 0, 24, 48 and 72 h time courses cold stress treatments. The shoot tissues of cold stress treated seedlings were used for quantitative proteomic analysis.

### Identification and quantitation of proteins with iTRAQ-based LC-MS/MS analysis

MS raw data were analyzed with ProteinPilot (V4.5) to identify and quantify proteins. As shown in Fig. 2A, a total of 89,976 MS/MS spectra were identified by iTRAQ-based LC-MS/MS analysis in time courses cold stress treated Kongyu131 shoot tissues. Among them, 29,601 peptides were found. At least one unique peptide identified for each confident protein. A total of 5082 unique proteins were identified by iTRAQ labeling from time courses cold-stressed Kongyu131 (Table S1).

**Fig 2.**
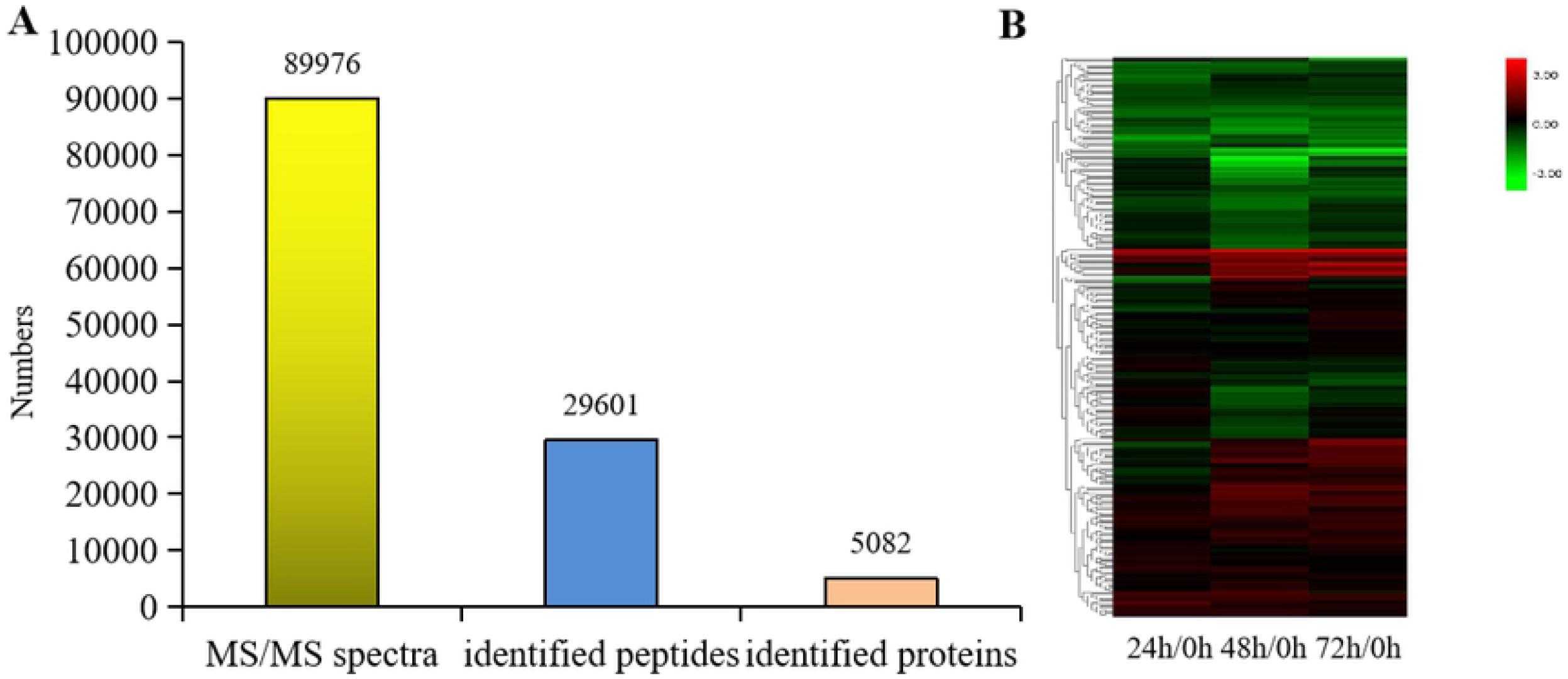
Basic information statistics of proteome and hierarchical clustering of DEPs identified during cold stress. A, Basic information statistics of proteome using iTRAQ analysis. MS/MS spectra are the secondary mass spectrums, and protein is identified by ProteinPilot (V4.5). B, Hierarchical clustering of quantified proteins based on LC-MS/MS data.

Through quantitative analysis with the software, 289 unique proteins were differently expressed with changes greater than 1.2-fold or smaller than 0.83-fold and ones with *p*-value smaller than 0.05 during cold stress treatment, stress and control samples formed two major clusters (Fig. 2B). After 24 h of cold treatment, 91 differentially expressed proteins (DEPs) (58 up- and 33 down-regulated) were found (Table 1). As the cold treatment time was increased, the number of slightly increased: 179 DEPs (86 up- and 93 down-regulated) at 48 h (Table 2) and 142 DEPs (98 up- and 44 down-regulated) at 72 h (Table 3). Fig. 3 showed the Venn diagram analysis of the DEPs at different time points. Overall, there were 289 unique DEPs (169 uniquely up-regulated and 125 uniquely down-regulated) during cold stress. Among the 169 up-regulated proteins, 11 proteins were found to be significantly up-regulated at three time points (Fig. 3A), and 11 proteins were found to be significantly down-regulated at three time points in 125 down-regulated proteins (Fig. 3B). Fifty-one proteins were found to be significantly up-regulated and 23 proteins were found to be significantly down-regulated at any two time points, respectively. Five proteins were found to be both significantly up- and down-regulated during cold stress treatments: 4 proteins up-regulated after 24 h of cold stress but down-regulated at 48 h time point, and one protein down-regulated after 48 h of cold stress but up-regulated at 72 h time point.

**Table 1.**
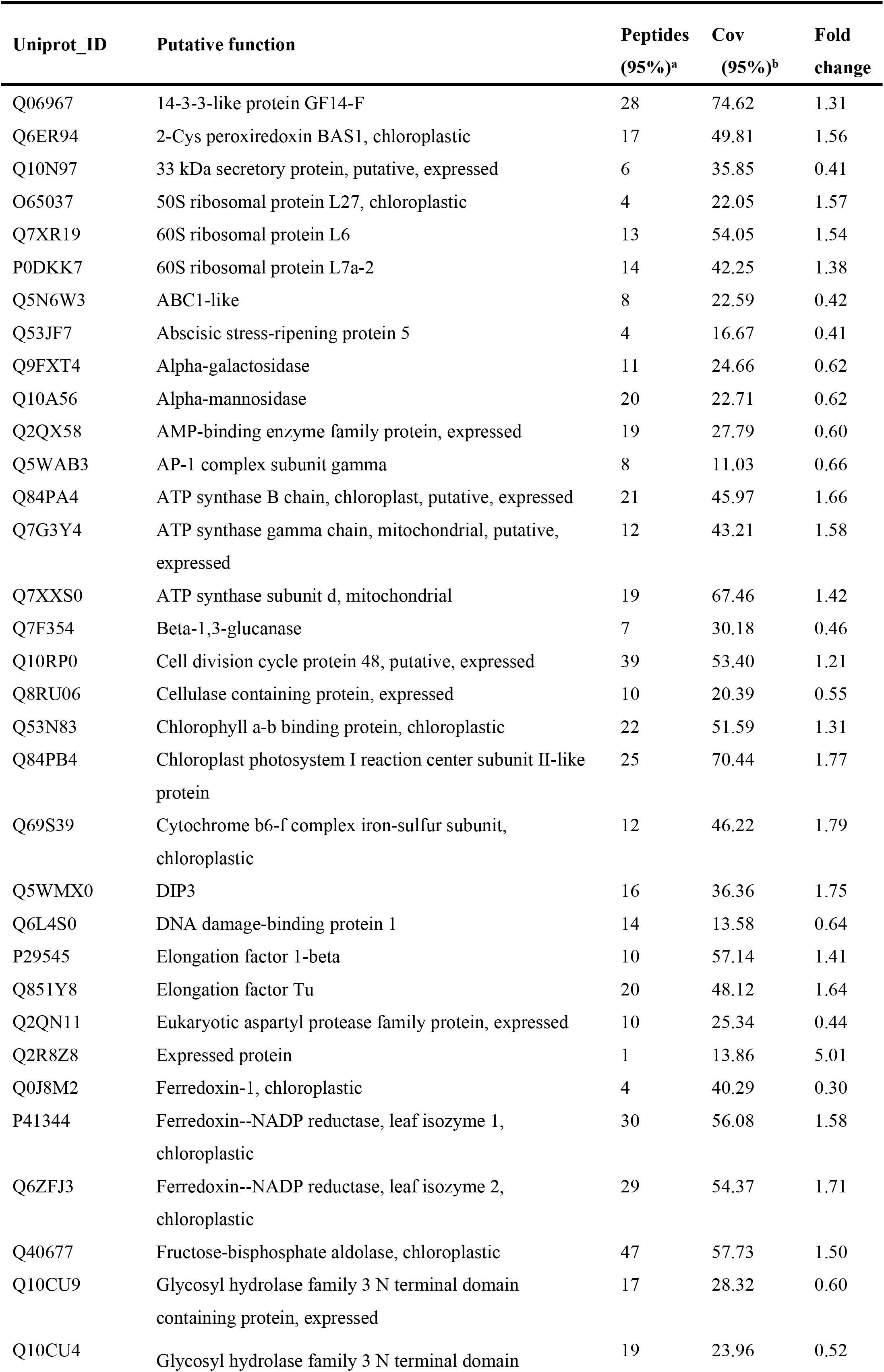
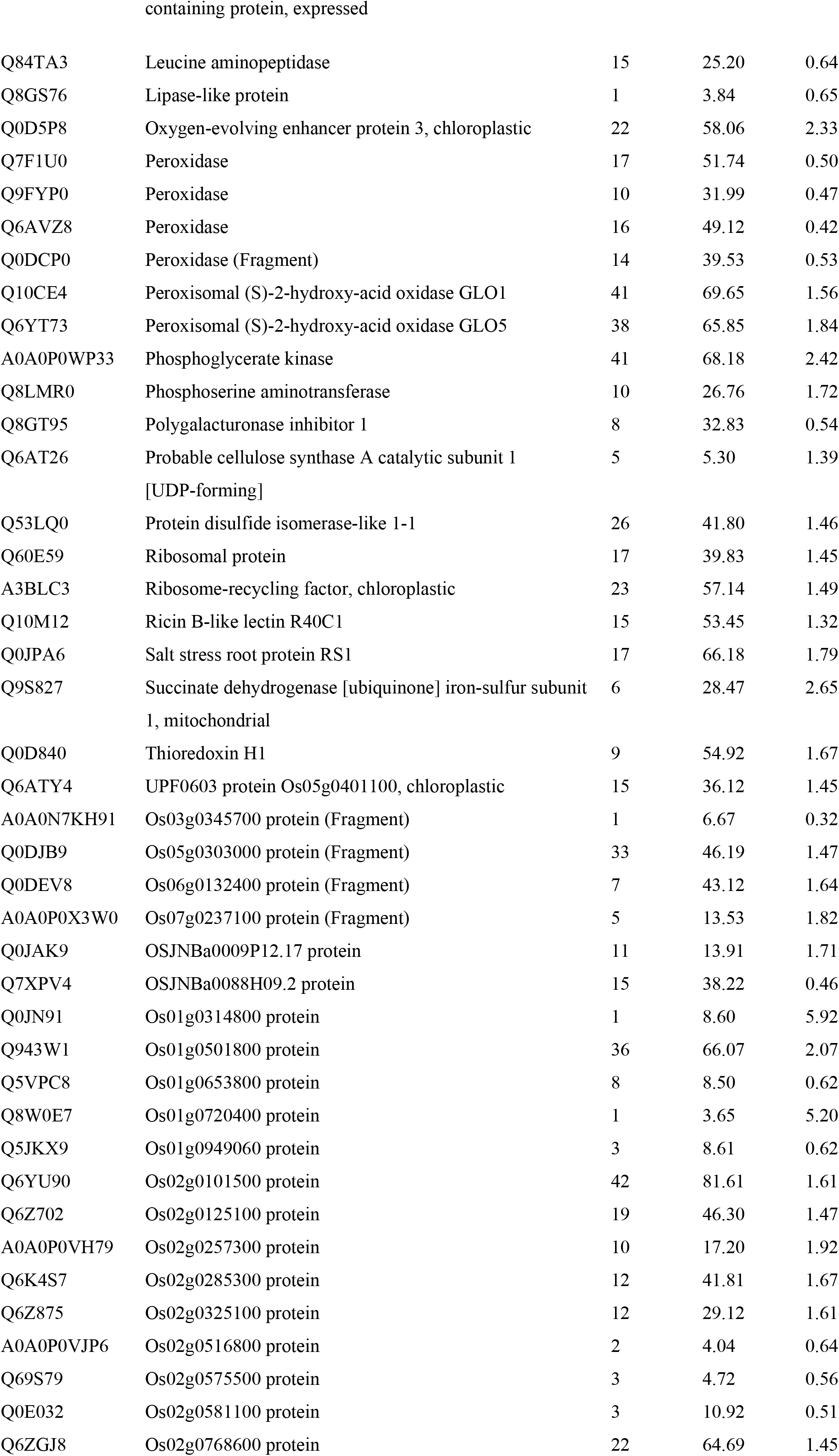
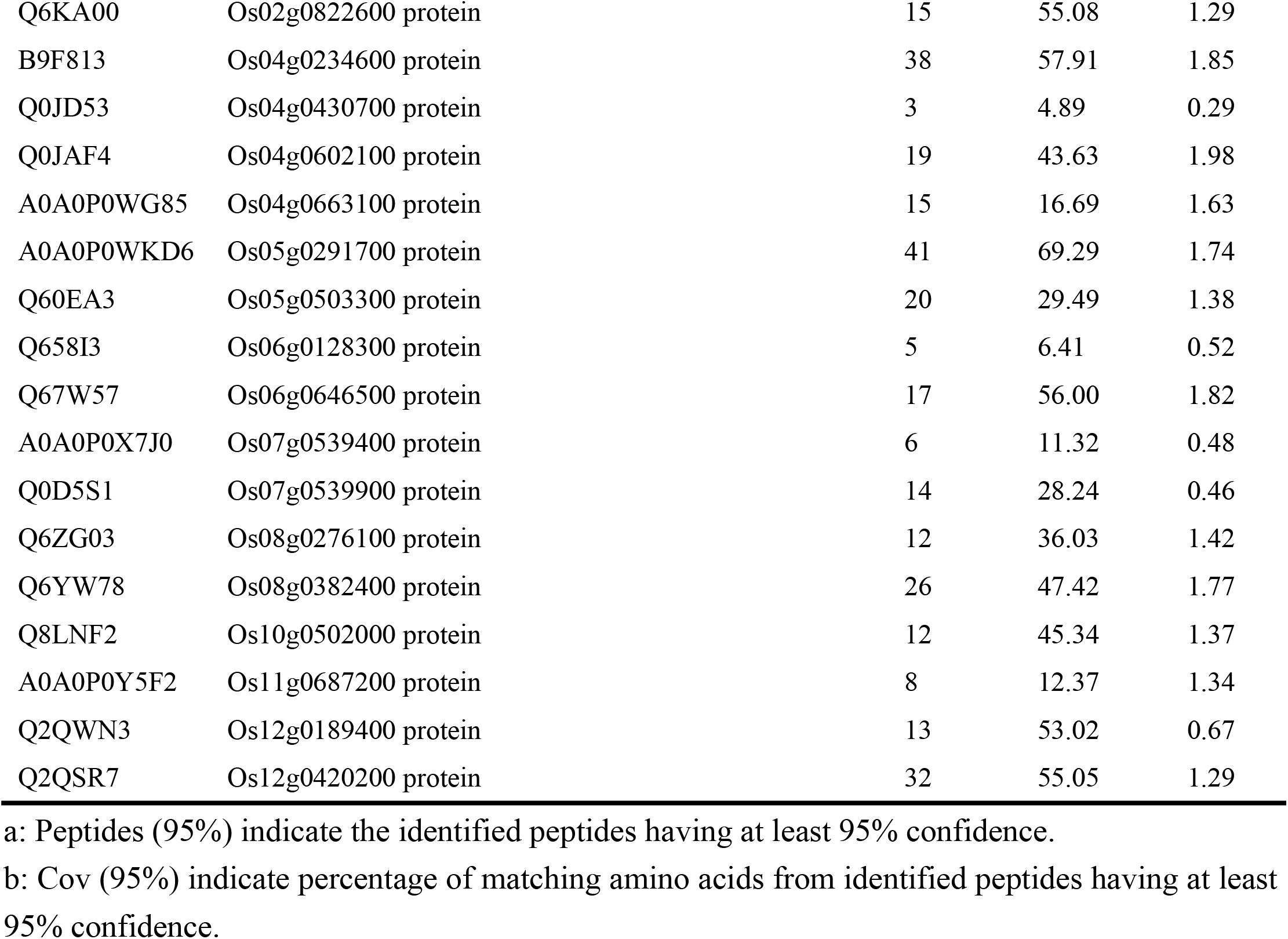
List of differentially expressed proteins after 24 h cold stress treatment.

**Table 2.**
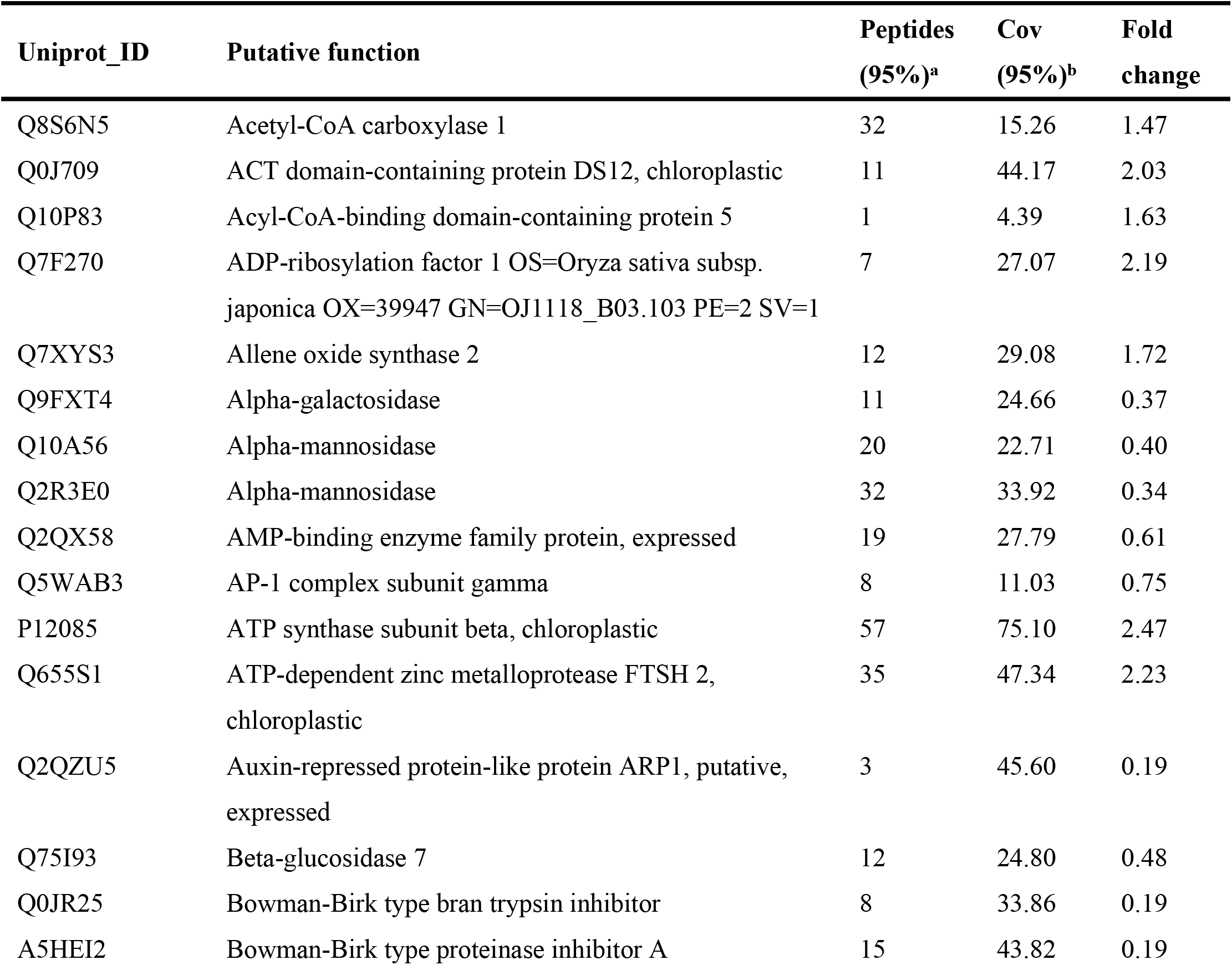
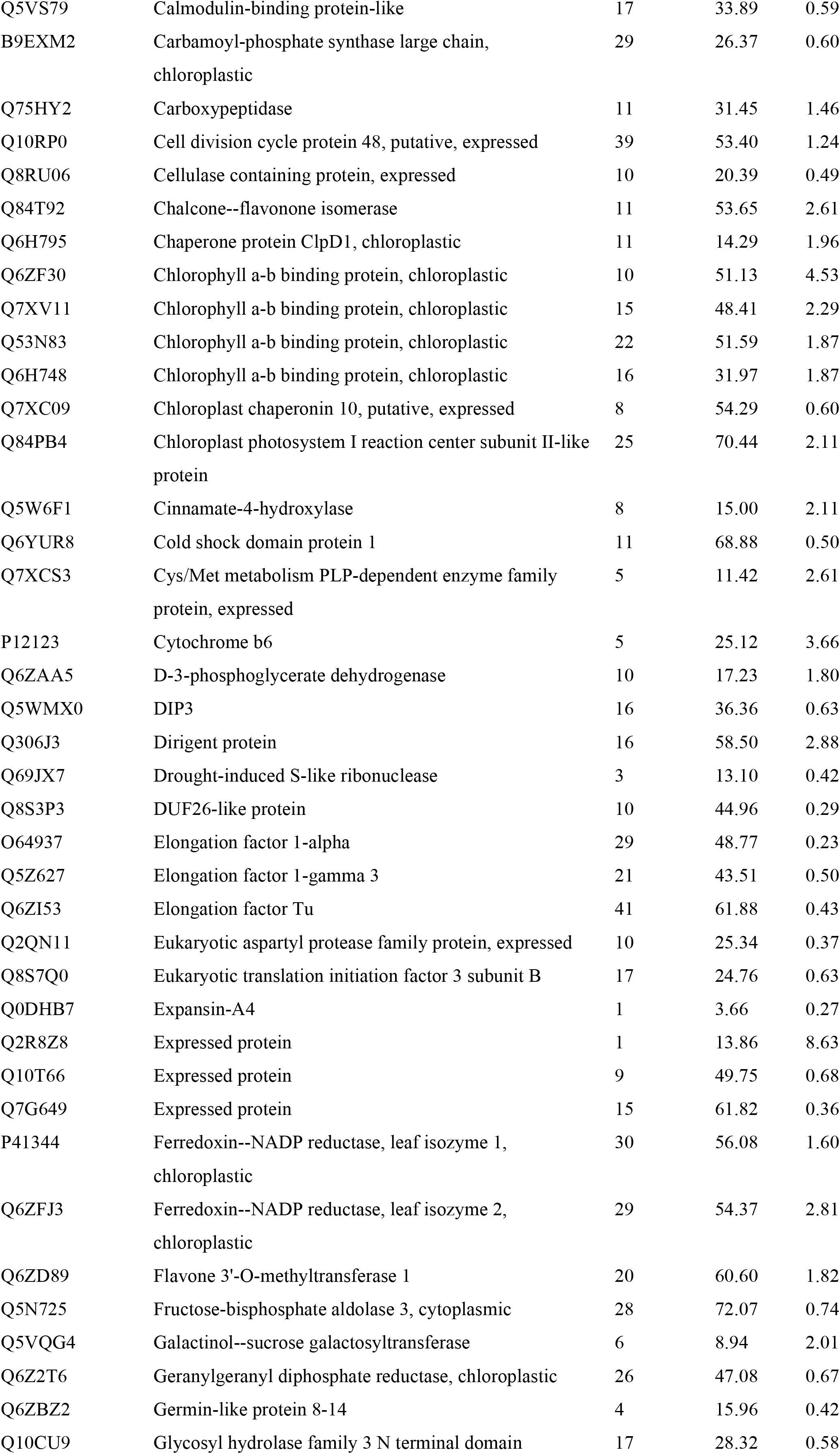
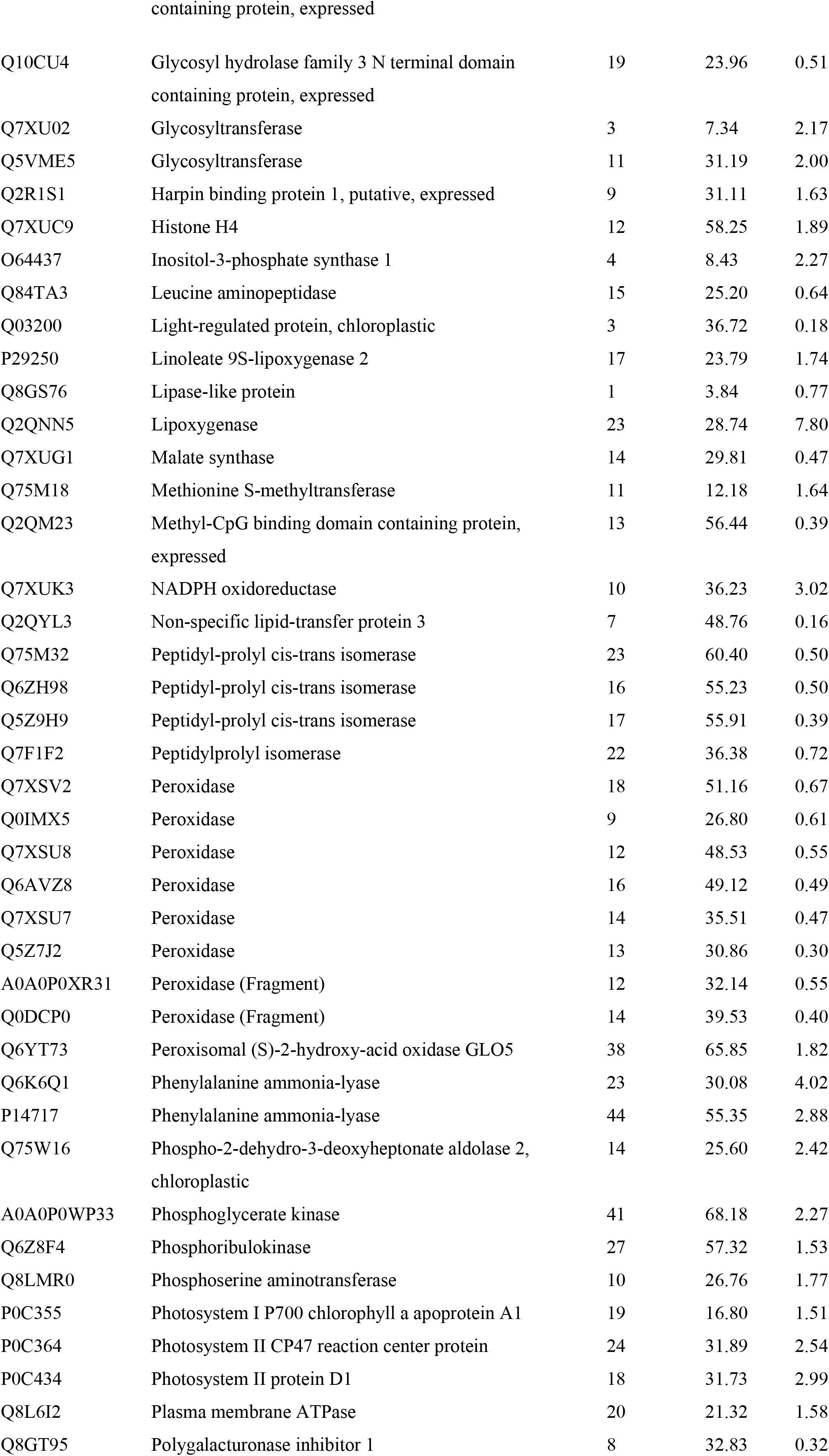
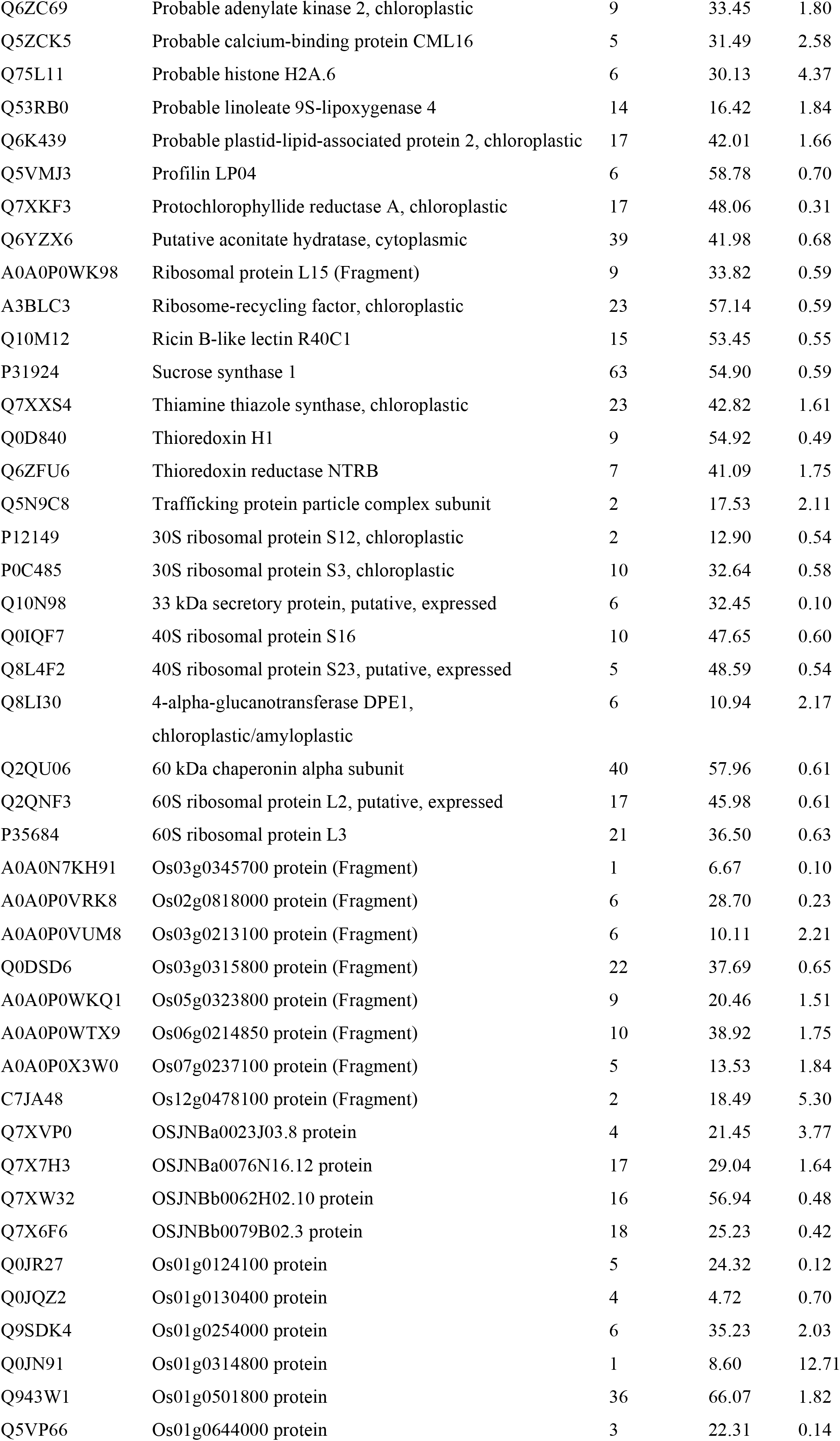
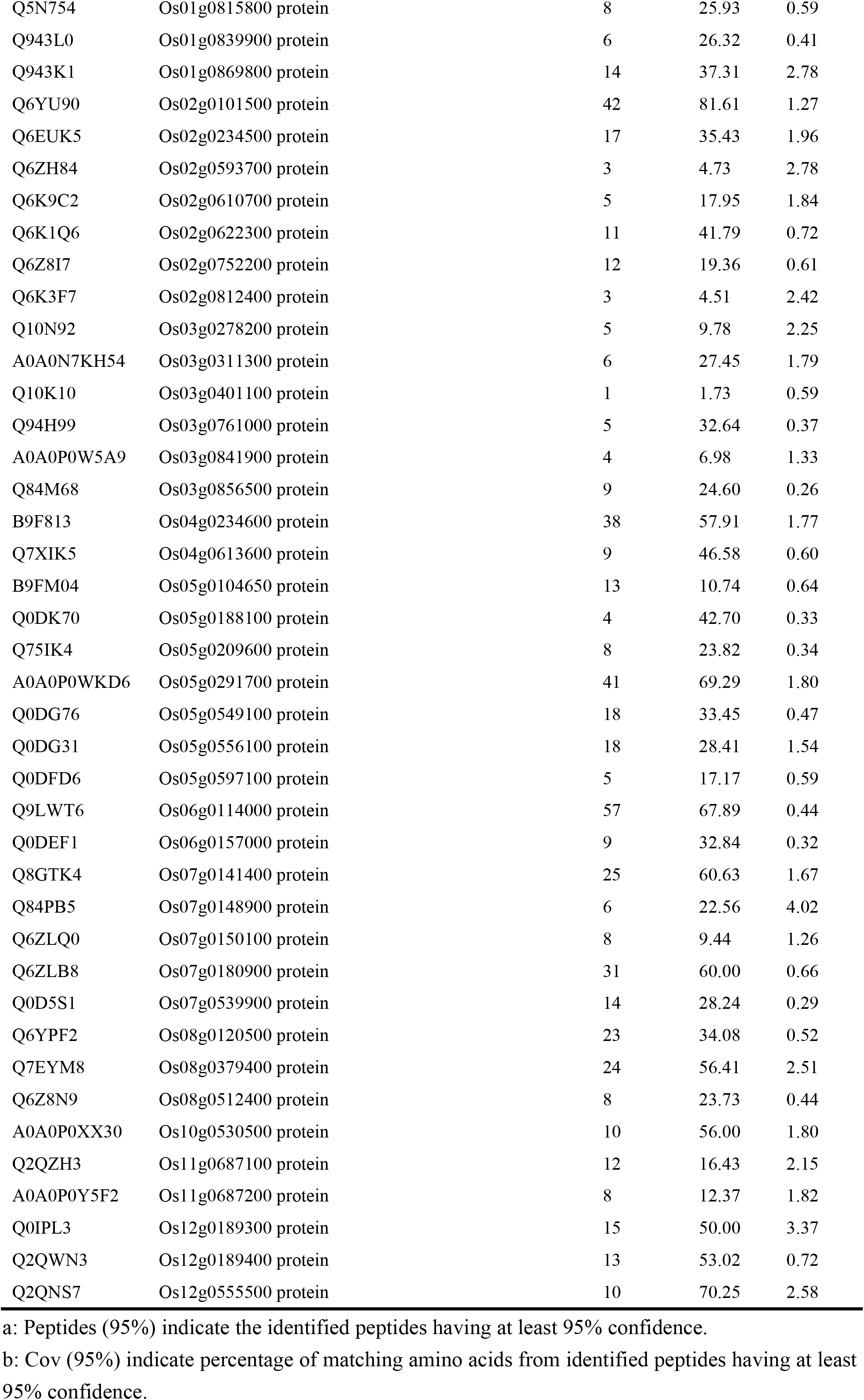
List of differentially expressed proteins after 48 h cold stress treatment.

**Table 3.** List of differentially expressed proteins after 72 h cold stress treatment.

| Uniprot_ID | Putative function | Peptides<br>(95%)<sup>a</sup> | Cov<br>(95%)<sup>b</sup> | Fold<br>change |
| --- | --- | --- | --- | --- |
| Q0J709 | ACT domain-containing protein DS12, chloroplastic | 11 | 44.17 | 2.03 |
| Q10P83 | Acyl-CoA-binding domain-containing protein 5 | 1 | 4.39 | 1.94 |
| A0A0P0Y1Y5 | Adenosylhomocysteinase (Fragment) | 30 | 55.15 | 1.27 |
| Q7XYS3 | Allene oxide synthase 2 | 12 | 29.08 | 1.72 |
| Q9FXT4 | Alpha-galactosidase | 11 | 24.66 | 0.43 |
| Q10A56 | Alpha-mannosidase | 20 | 22.71 | 0.67 |
| Q5WAB3 | AP-1 complex subunit gamma | 8 | 11.03 | 0.72 |
| P0C2Z6 | ATP synthase subunit alpha, chloroplastic | 36 | 51.68 | 1.20 |
| P0C522 | ATP synthase subunit alpha, mitochondrial | 31 | 53.24 | 1.71 |
| Q93VT8 | ATP-citrate synthase beta chain protein 1 | 25 | 46.71 | 1.43 |
| Q5Z974 | ATP-dependent zinc metalloprotease FTSH 1, chloroplastic | 34 | 40.38 | 1.32 |
| Q655S1 | ATP-dependent zinc metalloprotease FTSH 2, chloroplastic | 35 | 47.34 | 1.58 |
| Q0JR25 | Bowman-Birk type bran trypsin inhibitor | 8 | 33.86 | 0.39 |
| Q2QMX9 | Calcium-transporting ATPase 10, plasma membrane-type | 7 | 6.76 | 1.43 |
| B9EXM2 | Carbamoyl-phosphate synthase large chain, chloroplastic | 29 | 26.37 | 0.65 |
| Q75HY2 | Carboxypeptidase | 11 | 31.45 | 1.56 |
| Q10RP0 | Cell division cycle protein 48, putative, expressed | 39 | 53.40 | 1.79 |
| Q84T92 | Chalcone--flavonone isomerase | 11 | 53.65 | 2.51 |
| Q6H795 | Chaperone protein ClpD1, chloroplastic | 11 | 14.29 | 3.70 |
| Q6H748 | Chlorophyll a-b binding protein, chloroplastic | 16 | 31.97 | 2.00 |
| Q7XV11 | Chlorophyll a-b binding protein, chloroplastic | 15 | 48.41 | 2.13 |
| Q53N83 | Chlorophyll a-b binding protein, chloroplastic | 22 | 51.59 | 1.38 |
| Q6ZF30 | Chlorophyll a-b binding protein, chloroplastic | 10 | 51.13 | 3.08 |
| Q10HD0 | Chlorophyll a-b binding protein, chloroplastic | 16 | 45.25 | 2.73 |
| Q5W6F1 | Cinnamate-4-hydroxylase | 8 | 15.00 | 2.05 |
| Q6ZGV8 | Clustered mitochondria protein homolog | 8 | 7.36 | 1.53 |
| Q6YUR8 | Cold shock domain protein 1 | 11 | 68.88 | 0.48 |
| Q7XCS3 | Cys/Met metabolism PLP-dependent enzyme family protein,<br>expressed | 5 | 11.42 | 3.16 |
| Q7XKC8 | Dihydroorotate dehydrogenase (quinone), mitochondrial | 5 | 13.22 | 0.25 |
| Q306J3 | Dirigent protein | 16 | 58.50 | 1.82 |
| Q69JX7 | Drought-induced S-like ribonuclease | 3 | 13.10 | 0.49 |
| Q2QN11 | Eukaryotic aspartyl protease family protein, expressed | 10 | 25.34 | 0.43 |
| Q7G649 | Expressed protein | 15 | 61.82 | 0.52 |
| Q2R678 | Expressed protein | 11 | 27.67 | 1.54 |
| Q10T66 | Expressed protein | 9 | 49.75 | 0.51 |
| Q2R8Z8 | Expressed protein | 1 | 13.86 | 14.32 |
| P41344 | Ferredoxin--NADP reductase, leaf isozyme 1, chloroplastic | 30 | 56.08 | 1.77 |
| Q6ZFJ3 | Ferredoxin--NADP reductase, leaf isozyme 2, chloroplastic | 29 | 54.37 | 2.23 |
| Q6ZD89 | Flavone 3'-O-methyltransferase 1 | 20 | 60.60 | 1.96 |
| Q69V57 | Fructose-bisphosphate aldolase | 31 | 75.14 | 1.42 |
| Q40677 | Fructose-bisphosphate aldolase, chloroplastic | 47 | 57.73 | 1.47 |
| Q5VQG4 | Galactinol--sucrose galactosyltransferase OS=*Oryza sativa*<br>subsp. japonica OX=39947 GN=RFS PE=1 SV=1 | 6 | 8.94 | 2.49 |
| Q6Z2T6 | Geranylgeranyl diphosphate reductase, chloroplastic | 26 | 47.08 | 0.64 |
| P15280 | Glucose-1-phosphate adenylyltransferase small subunit 2,<br>chloroplastic/amyloplastic/cytosolic | 22 | 41.05 | 1.33 |
| P14656 | Glutamine synthetase cytosolic isozyme 1-1 | 9 | 33.99 | 1.74 |
| Q945W2 | Glutathione S-transferase GSTU6, putative, expressed | 6 | 28.81 | 2.15 |
| Q8H8D6 | Glutathione S-transferase, N-terminal domain containing<br>protein, expressed | 16 | 46.39 | 0.62 |
| Q0J8A4 | Glyceraldehyde-3-phosphate dehydrogenase 1, cytosolic | 28 | 62.02 | 2.81 |
| A3C6G9 | Glycine cleavage system H protein, mitochondrial | 8 | 67.68 | 0.41 |
| Q10CU9 | Glycosyl hydrolase family 3 N terminal domain containing<br>protein, expressed | 17 | 28.32 | 0.60 |
| Q7XU02 | Glycosyltransferase | 3 | 7.34 | 2.65 |
| Q5VME5 | Glycosyltransferase | 11 | 31.19 | 1.82 |
| Q7XUC9 | Histone H4 | 12 | 58.25 | 2.96 |
| Q851P9 | Histone-like protein | 6 | 19.11 | 3.73 |
| O64437 | Inositol-3-phosphate synthase 1 | 4 | 8.43 | 2.40 |
| Q84TA3 | Leucine aminopeptidase | 15 | 25.20 | 0.65 |
| Q6K669 | Leucine aminopeptidase 2, chloroplastic | 27 | 39.63 | 1.29 |
| P38419 | Lipoxygenase 7, chloroplastic | 16 | 18.94 | 1.54 |
| Q7XZW5 | Malate dehydrogenase | 34 | 85.31 | 1.64 |
| Q2QM23 | Methyl-CpG binding domain containing protein, expressed | 13 | 56.44 | 0.52 |
| Q7XUK3 | NADPH oxidoreductase | 10 | 36.23 | 3.70 |
| Q0D5P8 | Oxygen-evolving enhancer protein 3, chloroplastic | 22 | 58.06 | 1.25 |
| Q5JMS4 | Peroxidase | 18 | 50.40 | 1.79 |
| Q6AVZ8 | Peroxidase | 16 | 49.12 | 0.44 |
| Q9FYP0 | Peroxidase | 10 | 31.99 | 0.48 |
| Q0JM38 | Peroxidase | 8 | 25.59 | 0.27 |
| Q5Z7J2 | Peroxidase | 13 | 30.86 | 0.47 |
| Q0DCP0 | Peroxidase (Fragment) | 14 | 39.53 | 0.50 |
| A0A0P0XR31 | Peroxidase (Fragment) | 12 | 32.14 | 0.57 |
| Q6YT73 | Peroxisomal (S)-2-hydroxy-acid oxidase GLO5 | 38 | 65.85 | 2.01 |
| P14717 | Phenylalanine ammonia-lyase | 44 | 55.35 | 2.70 |
| Q6K6Q1 | Phenylalanine ammonia-lyase | 23 | 30.08 | 3.98 |
| Q0DZE0 | Phenylalanine ammonia-lyase | 22 | 27.91 | 5.86 |
| Q75W16 | Phospho-2-dehydro-3-deoxyheptonate aldolase 2,<br>chloroplastic | 14 | 25.60 | 2.49 |
| A0A0P0WP33 | Phosphoglycerate kinase | 41 | 68.18 | 1.92 |
| P0C355 | Photosystem I P700 chlorophyll a apoprotein A1 | 19 | 16.80 | 1.56 |
| P0C358 | Photosystem I P700 chlorophyll a apoprotein A2 | 20 | 20.84 | 2.56 |
| P0C364 | Photosystem II CP47 reaction center protein | 24 | 31.89 | 2.75 |
| Q8GT95 | Polygalacturonase inhibitor 1 | 8 | 32.83 | 0.43 |
| Q5ZCK5 | Probable calcium-binding protein CML16 | 5 | 31.49 | 2.05 |
| Q75L11 | Probable histone H2A.6 | 6 | 30.13 | 4.41 |
| Q6K439 | Probable plastid-lipid-associated protein 2, chloroplastic | 17 | 42.01 | 1.79 |
| Q0D5W6 | Protein translation factor SUI1 homolog | 6 | 43.48 | 2.68 |
| Q6ZHE5 | Putative D-cysteine desulfhydrase 1, mitochondrial | 13 | 34.98 | 1.34 |
| Q7XKB5 | Pyruvate kinase | 18 | 32.09 | 1.22 |
| P31924 | Sucrose synthase 1 | 63 | 54.90 | 1.33 |
| Q93X08 | UTP--glucose-1-phosphate uridylyltransferase | 35 | 60.55 | 1.39 |
| Q06967 | 14-3-3-like protein GF14-F | 28 | 74.62 | 1.43 |
| Q6ER94 | 2-Cys peroxiredoxin BAS1, chloroplastic | 17 | 49.81 | 1.45 |
| P0C488 | 30S ribosomal protein S4, chloroplastic | 11 | 42.29 | 2.03 |
| Q8LI30 | 4-alpha-glucanotransferase DPE1, chloroplastic/amyloplastic | 6 | 10.94 | 2.09 |
| Q2QLY5 | 5-methyltetrahydropteroyltriglutamate--homocysteine methyltransferase 1 | 40 | 43.99 | 2.03 |
| P49210 | 60S ribosomal protein L9 | 11 | 55.26 | 1.54 |
| Q2R480 | 6-phosphogluconate dehydrogenase, decarboxylating 2, chloroplastic | 17 | 37.20 | 1.43 |
| A0A0N7KH91 | Os03g0345700 protein (Fragment) | 1 | 6.67 | 0.12 |
| A0A0P0VRK8 | Os02g0818000 protein (Fragment) | 6 | 28.70 | 0.29 |
| A0A0P0VTG7 | Os03g0165300 protein (Fragment) | 4 | 8.65 | 0.18 |
| A0A0P0VUM8 | Os03g0213100 protein (Fragment) | 6 | 10.11 | 2.38 |
| Q0DEU8 | Os06g0133800 protein (Fragment) | 52 | 64.45 | 1.50 |
| A0A0P0Y445 | Os11g0602500 protein (Fragment) | 13 | 37.32 | 1.71 |
| C7JA48 | Os12g0478100 protein (Fragment) | 2 | 18.49 | 11.91 |
| Q7XPV4 | OSJNBa0088H09.2 protein | 15 | 38.22 | 0.68 |
| Q7XVF8 | OSJNBb0118P14.7 protein | 5 | 15.38 | 2.00 |
| Q94EG1 | Os01g0178000 protein | 7 | 24.37 | 0.69 |
| Q5VR43 | Os01g0180300 protein | 51 | 55.48 | 0.61 |
| Q9SDK4 | Os01g0254000 protein | 6 | 35.23 | 1.57 |
| Q9FP25 | Os01g0303000 protein | 14 | 62.90 | 0.44 |
| Q0JN91 | Os01g0314800 protein | 1 | 8.60 | 8.32 |
| Q6YU90 | Os02g0101500 protein | 42 | 81.61 | 1.58 |
| Q6YUV4 | Os02g0189000 protein | 6 | 32.88 | 0.51 |
| Q6EUK5 | Os02g0234500 protein | 17 | 35.43 | 1.67 |
| Q6ZH84 | Os02g0593700 protein | 3 | 4.73 | 4.83 |
| Q6Z8I7 | Os02g0752200 protein | 12 | 19.36 | 0.62 |
| Q0DWG3 | Os02g0816300 protein | 4 | 3.93 | 0.40 |
| Q10RI4 | Os03g0158500 protein | 5 | 8.77 | 2.17 |
| Q10N92 | Os03g0278200 protein | 5 | 9.78 | 2.25 |
| Q9AUQ4 | Os03g0712700 protein | 32 | 51.03 | 1.42 |
| Q8S7H8 | Os03g0778100 protein | 11 | 37.29 | 2.29 |
| B9F813 | Os04g0234600 protein | 38 | 57.91 | 1.89 |
| Q0JAF4 | Os04g0602100 protein | 19 | 43.63 | 1.77 |
| Q75IR7 | Os05g0163000 protein | 15 | 26.73 | 1.46 |
| Q0DK70 | Os05g0188100 protein | 4 | 42.70 | 0.42 |
| Q75IK4 | Os05g0209600 protein | 8 | 23.82 | 0.57 |
| Q6F322 | Os05g0490700 protein | 9 | 13.84 | 1.33 |
| Q0DG76 | Os05g0549100 protein | 18 | 33.45 | 0.51 |
| Q9SNN5 | Os06g0130800 protein | 8 | 30.83 | 0.69 |
| Q7XXQ8 | Os06g0232000 protein | 6 | 16.80 | 0.36 |
| Q69WH2 | Os06g0332800 protein | 6 | 44.27 | 0.43 |
| Q93W07 | Os06g0568200 protein | 23 | 50.82 | 1.24 |
| Q67W57 | Os06g0646500 protein | 17 | 56.00 | 2.03 |
| Q5YM05 | Os06g0720400 protein | 10 | 22.75 | 1.58 |
| Q8GTK4 | Os07g0141400 protein | 25 | 60.63 | 1.57 |
| Q84PB5 | Os07g0148900 protein | 6 | 22.56 | 4.66 |
| A0A0P0X7J0 | Os07g0539400 protein | 6 | 11.32 | 0.40 |
| Q0D5S1 | Os07g0539900 protein | 14 | 28.24 | 0.63 |
| Q0D3Z0 | Os07g0658300 protein | 1 | 1.03 | 0.72 |
| Q6ZFI6 | Os08g0502700 protein | 29 | 58.21 | 1.82 |
| Q650W6 | Os09g0565200 protein | 13 | 36.65 | 0.54 |
| Q2QZH3 | Os11g0687100 protein | 12 | 16.43 | 1.67 |
| A0A0P0Y5F2 | Os11g0687200 protein | 8 | 12.37 | 1.72 |
| Q0IPL3 | Os12g0189300 protein | 15 | 50.00 | 2.51 |
| Q2QSR7 | Os12g0420200 protein | 32 | 55.05 | 1.33 |

a: Peptides (95%) indicate the identified peptides having at least 95% confidence.
b: Cov (95%) indicate percentage of matching amino acids from identified peptides having at least 95% confidence.

**Fig 3.**
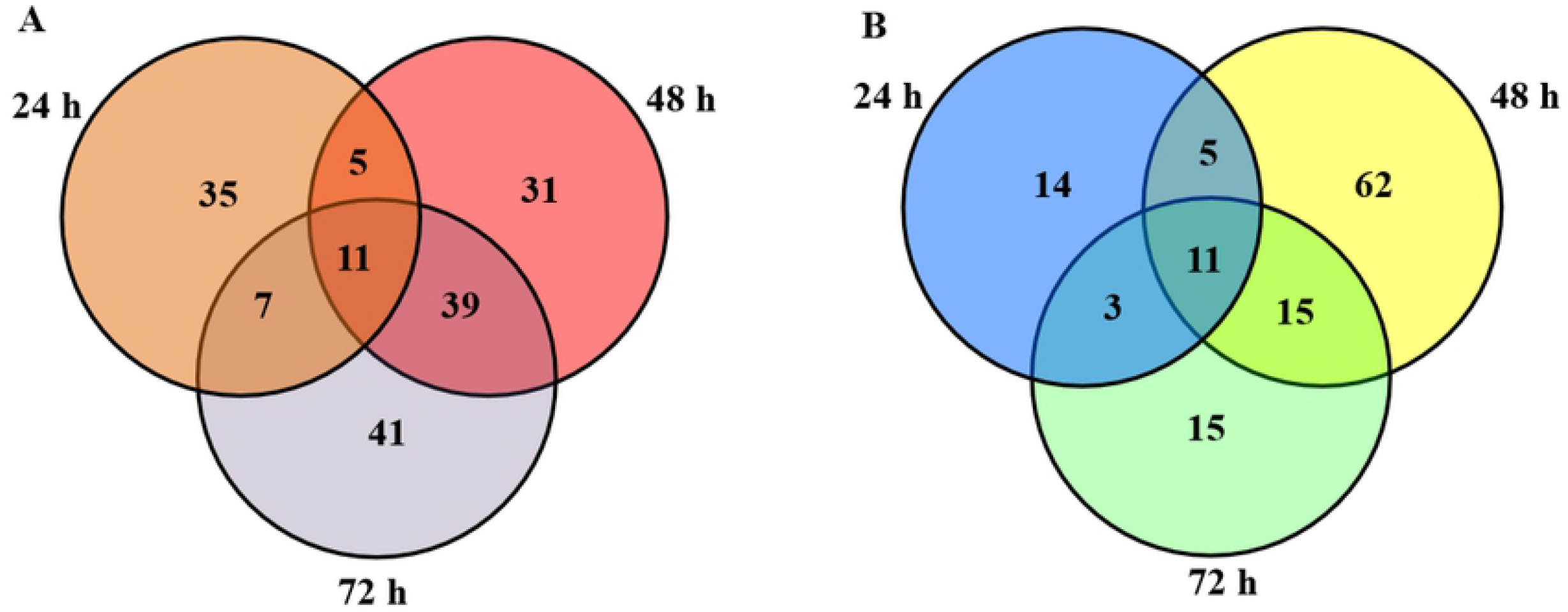
Venn diagram analysis of the differentially expressed proteins in rice at each time point cold stress treatment. The numbers of the differentially expressed proteins identified after 24 h, 48 h and 72 h cold treatment are shown in the different segments. A, The up-regulated proteins. B, The down-regulated proteins.

### GO analysis of DEPs

To further understand the functions of DEPs, GO analysis was performed. 253 protein IDs of 289 unique DEPs were assigned functions in the GO analysis. The DEPs were significantly enriched in 13/14/11 biological processes at 24/48/72 h cold stress treatment, 10/10/8 cellular components at 24/48/72 h cold stress treatment, and 7/8/6 molecular function subgroups at 24/48/72 h cold stress treatment. The metabolic process, cellular process and response to stimulus groups were prominent in the biological process subgroup, indication that the metabolic processes are more quickly affected under cold stress (Fig. 4A). The cell part, organelle, organelle part, membrane part and protein-containing complex groups were highly localized within the cellular component subgroup (Fig. 4B). Among the DEPs, the enriched GO terms concerning molecular function showed that DEPs were mainly associated with catalytic activity and binding, followed by the structural molecule activity, antioxidant activity and molecular function regulator (Fig. 4C).

**Fig 4.**
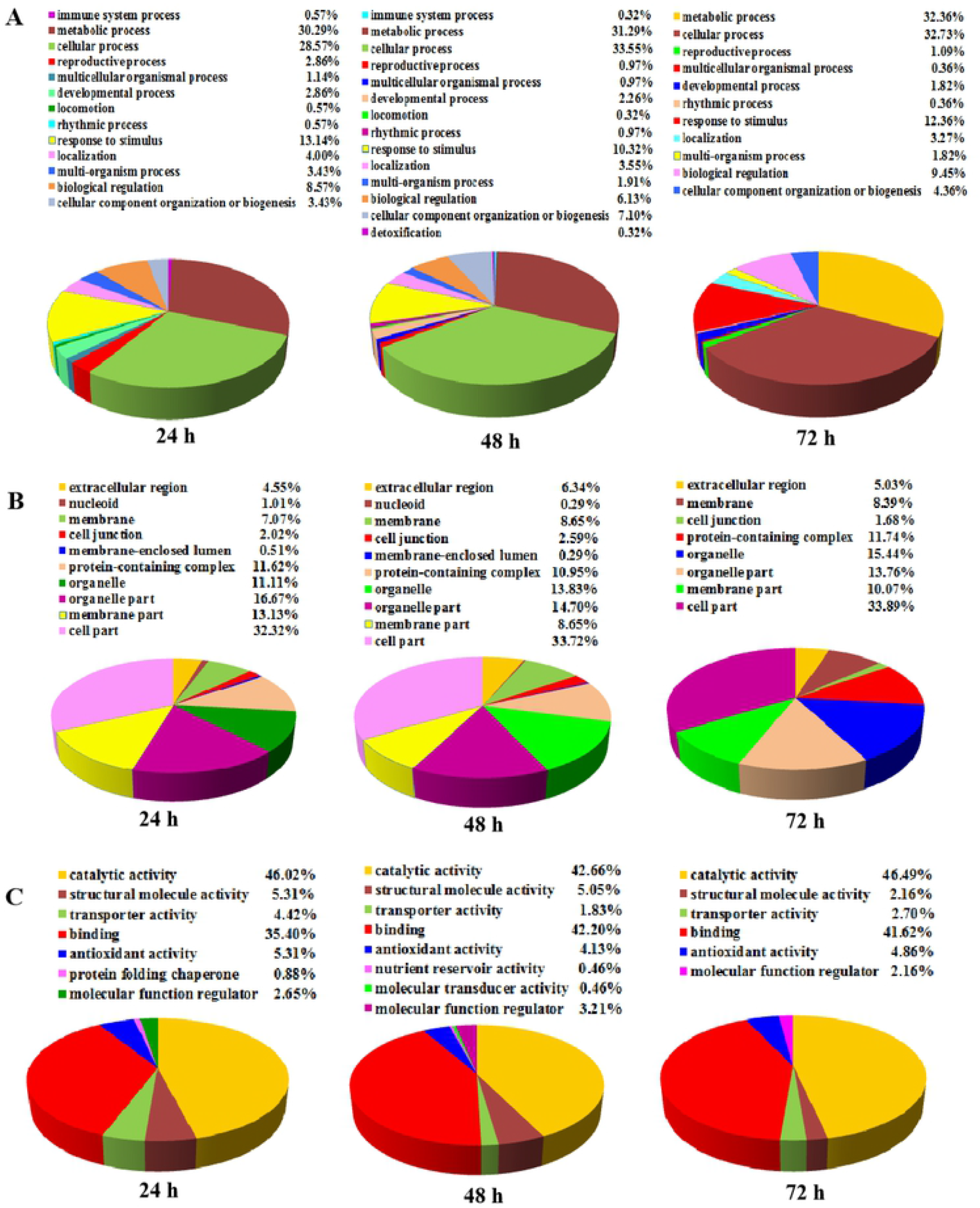
Bioinformatics analysis of DEPs in rice after 24 h, 48 h and 72 h cold stress treatments (ratio > 1.2 or < 0.83). A, Biological process. B, Cellular component. C, Molecular function.

### Pathway enrichment analysis of DEPs

DEPs of 24 h, 48 h and 72 h cold treatment were mapped to the reference pathway in the KEGG database for functional analysis. The metabolic pathways, photosynthesis, phenylpropanoid biosynthesis, carbon metabolism and carbon fixation in photosynthetic organisms were significantly enriched in three time points cold stress treatment (Fig. 5). There were some different pathways enriched in different cold tress time points, for example, oxidative phosphorylation, glucosinolate biosynthesis, vitamin B6 metabolism and so on were specific enriched at 24 h cold stress treatment (Fig. 5A); linoleic acid metabolism, cyanoamino metabolism, thiamine metabolism and zeatin biosynthesis were specific enriched at 48 h cold stress treatment (Fig. 5B); cysteine and methionine metabolism, galactose metabolism pentose phosphate pathway and starch/sucrose metabolism were specific enriched at 72 h cold stress treatment (Fig. 5C). With the cold stress time increase, enriched pathways more and more stable, there were ten of the same pathways both at 48 h and 72 h cold stress time points (Fig. 5).

**Fig 5.**
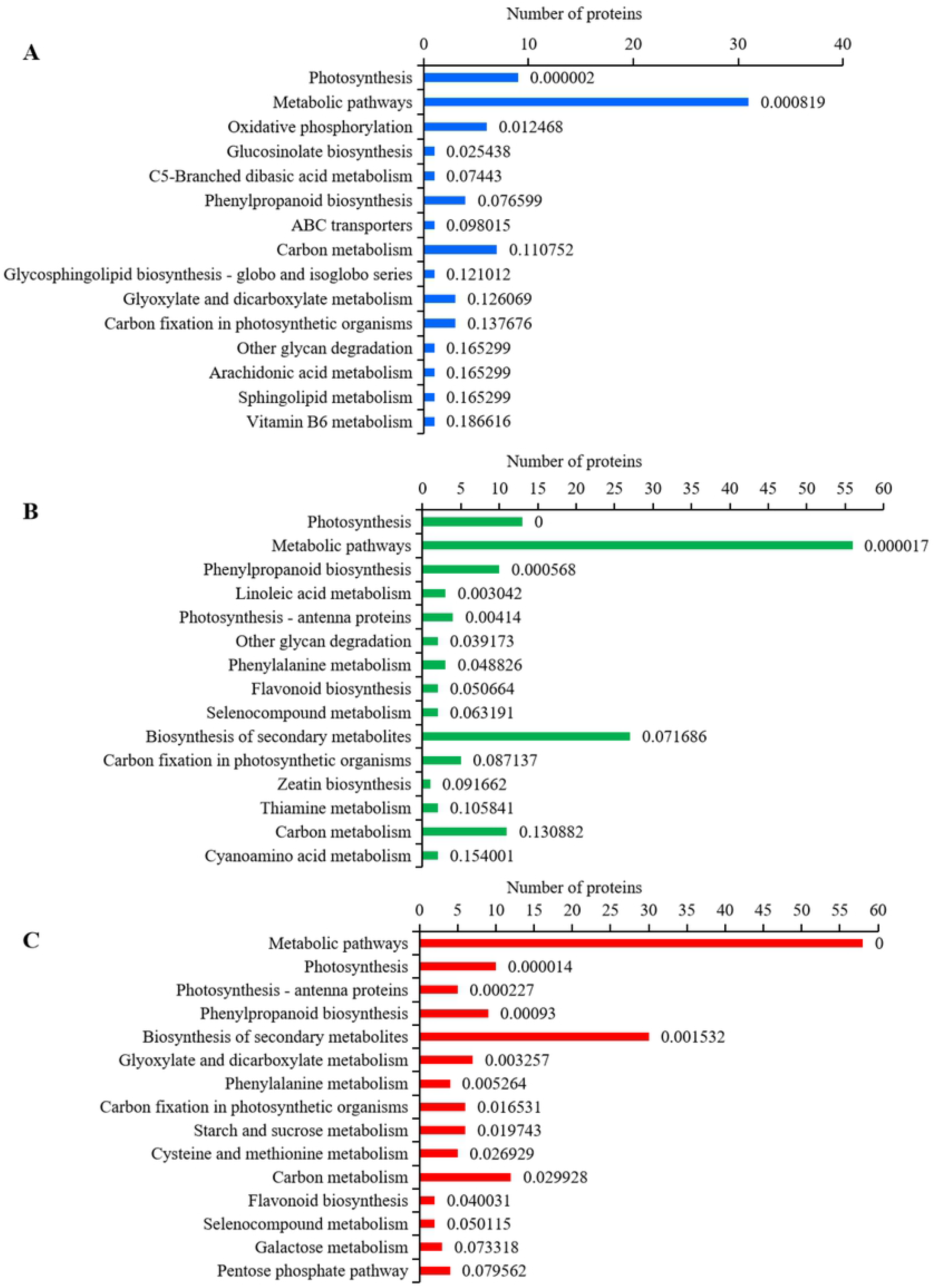
KEGG pathway analysis of DEPs in rice with cold stress treatments (ratio>1.2 or < 0.83). A, DEPs of 24 h cold treatment. B, DEPs of 48 h cold treatment. C, DEPs of 72 h cold treatment.

### Protein-protein interaction analysis of DEPs

The protein-protein interaction networks of DEPs from three time points cold stress treatment rice contain biological processes, cellular components and molecular function were constructed basing on the Search Tool for the Retrieval of Interacting Genes/Proteins 11.0 (STRING 11.0) database. By removing unconnected proteins, the resulting network of 24 h cold response proteins contained 64 protein nodes and 360 edges (Fig. 6A), the resulting network of 48 h cold response proteins contained 121 protein nodes and 546 edges (Fig. 6B), and the resulting network of 72 h cold response proteins contained 90 protein nodes and 402 edges (Fig. 6C). In biological processes, DEPs that function in metabolic process, cellular process, response to stimulus and biological regulation are highly up-regulated during cold stress; in terms of cellular component, DEPs that function in extracellular region, membrane, protein-containing complex and cell part are up-regulated during cold stress; in terms of molecular function, DEPs that function in catalytic activity, binding and molecular function regulator are highly up-regulated during cold tress.

**Fig 6.**
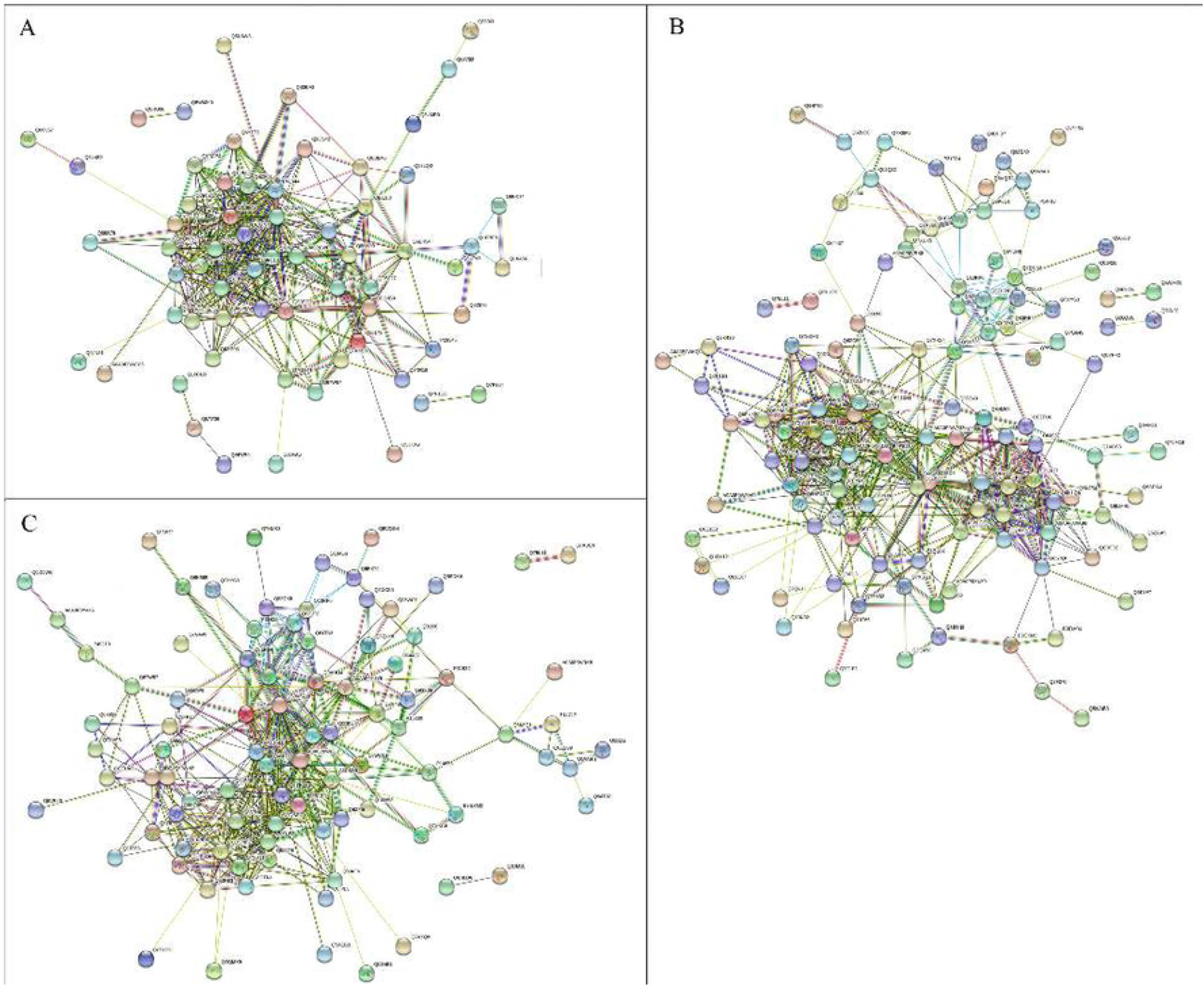
The protein-protein interaction network of cold response proteins generated using the STRING database. Medium confidence (STRING score = 0.4) was set in the network analysis. The edges represent predicted protein-protein associations. A, DEPs of 24 h cold stress treatment. B, DEPs of 48 h cold stress treatment. C, DEPs of 72 h cold stress treatment. Lake blue lines represent from curated databases, pink lines represent experimentally determined, green lines represent gene neighborhood, red lines represent gene fusions, dark blue lines represent gene co-occurrence, yellow lines represent textmining, black lines represent co-expression, sky blue lines represent protein homology.

### Validation of iTRAQ data on selected candidates by Q-PCR and western blot

19 up-regulated DEPs and 6 down-regulated DEPs were selected for Q-PCR analysis, to validate the relationship of expression profiles between mRNAs and proteins level. We compare the transcription levels of 24 h, 48 h and 72 h cold stress treatments with the iTRAQ data. As shown in Fig. 7, Q-PCR data indicated that the mRNA levels of 19 up-regulated DEPs increased under cold stress, the regulation trends of four DEPs (Q7XUK3, Q6ZH84, Q6ZFJ3, A0A0P0WP33) at three time points cold treatment were consistent with the iTRAQ quantification data. The mRNA levels of four out of six down-regulated DEPs decreased under cold stress, and the regulation trends of two DEPs (A0A0N7KH91and Q9FXT4) at three time points cold treatment were consistent with the iTRAQ quantification data (Fig. 7).

**Fig 7.**
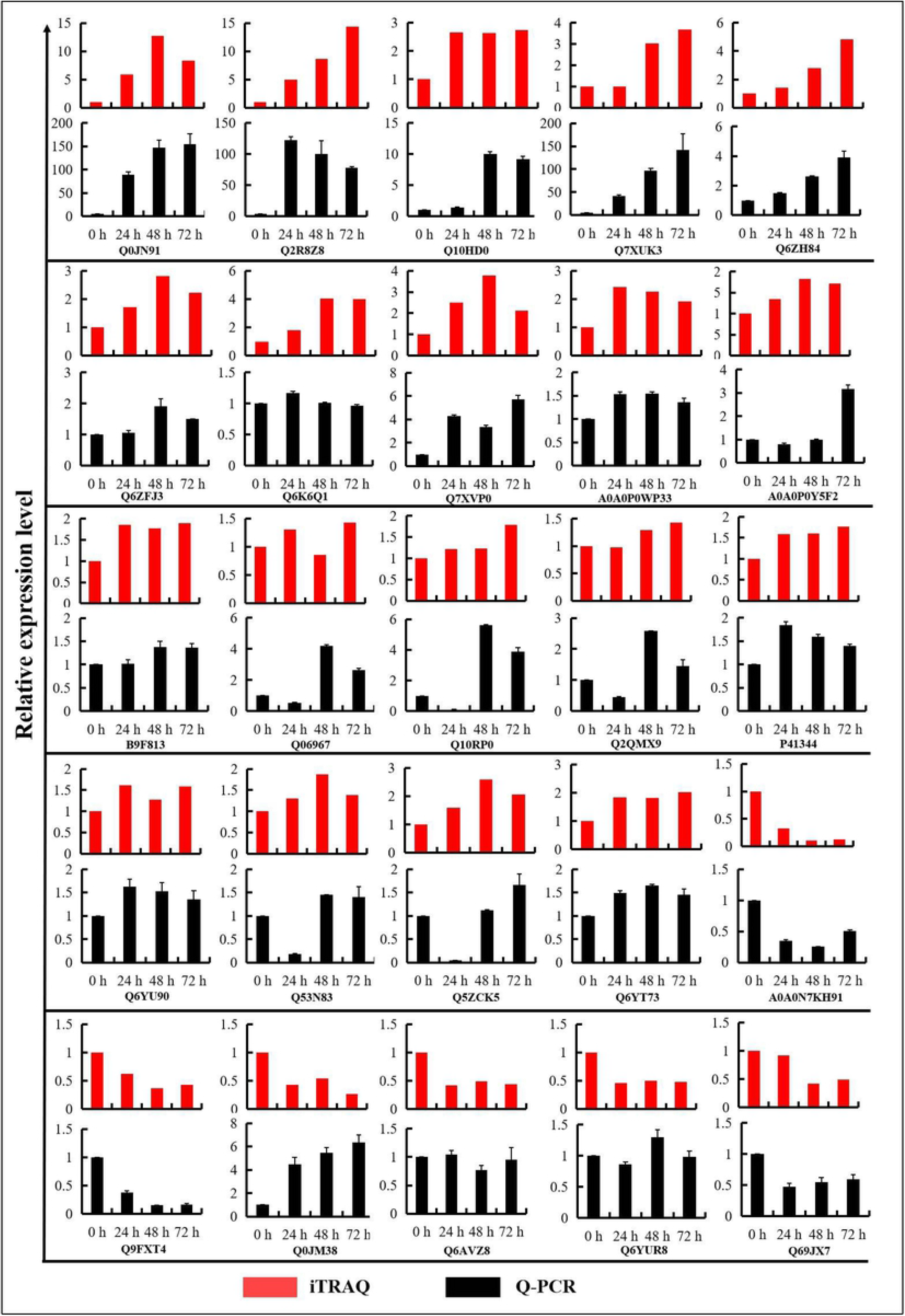
Comparative analysis of the protein and mRNA profiles of 25 representative DEPs. The X-axis represents the time points in the cold treatments. The Y-axis indicates the normalized relative protein and mRNA levels. The red and black columns represent the patterns of protein and mRNA expression in Kongyu131, respectively.

To further validate the protein regulation levels of DEPs identified by the iTRAQ labeling analysis, one down-regulated protein A0A0N7KH91 was selected for confirmation by western blot analysis with specific peptide antibody raised against the protein, and β-actin antibody as control. Fig. 8 showed that A0A0N7KH91 protein also down-regulated during cold stress treatment from 24 h to 72 h. This result further confirmed the iTRAQ labeling analysis data.

**Fig 8.**
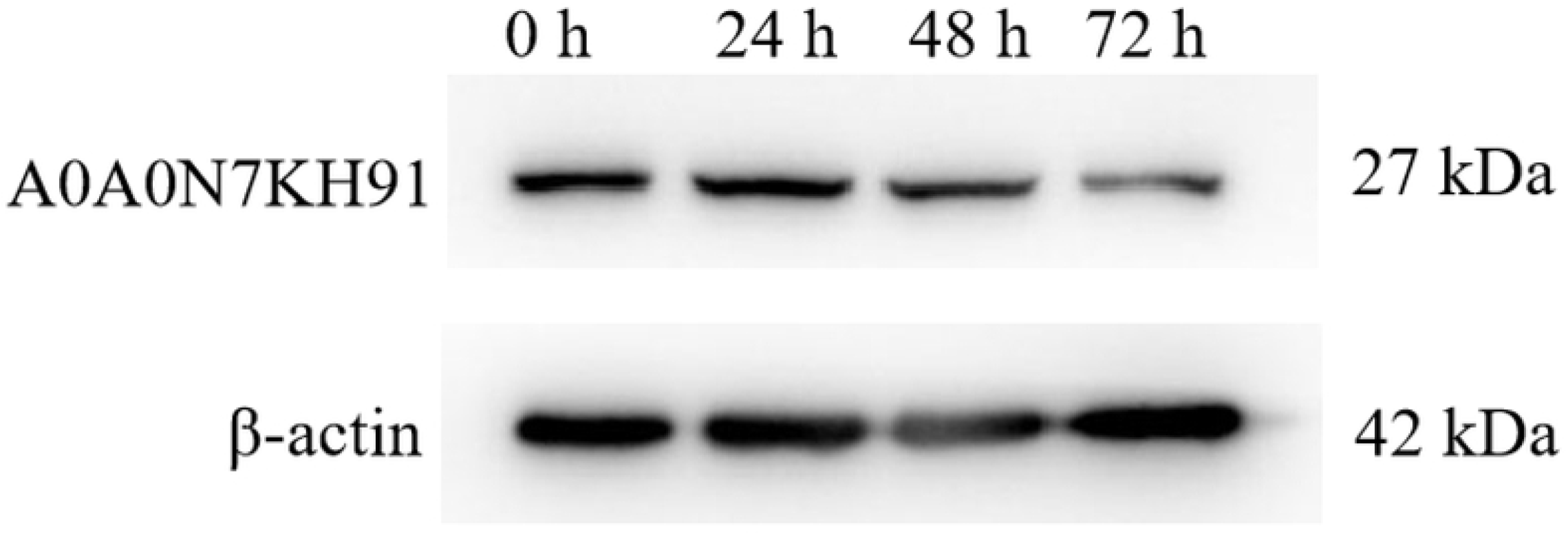
Western blot analysis of time courses cold-treated rice cultivar Kongyu131. Equal amount of 50 μg of total protein from different samples was used for western blot analysis with the enhanced chemiluminescence (ECL) approach. The specific antibodies against A0A0N7KH91 protein (1:500) and β-actin (1:5000) were used to detect the corresponding protein expressions.

## Discussion

In this work, we performed quantitative proteomic analysis of *japonica* rice seedlings subjected to time course cold stress treatments to obtain the dynamic proteins expression patterns responsive to low temperature. Using iTRAQ labeling couple with LC-MS/MS analysis approach, 5802 proteins were identified and were used for quantification from the rice tissues. As a result, we found 91/179/142 cold responsive proteins at 24/48/72 h cold stress treatments with the fold change > 1.2 or < 0.83 with a p-value < 0.05 for the differentially regulated proteins (Table 1-3), and the number of cold responsive proteins increased when the treatment time increased. And this result consistent with the data of previous studies of quantitative proteomic analysis of *indica* rice with time course cold stress treatment [29].

More and more proteomics studies on rice cold stress treatment to explore more cold response proteins for understanding plant cold-tolerance mechanism. Some proteins that were identified previously to be cold response proteins are further confirmed in our study using quantitative proteomic method. These proteins mainly include: sucrose synthase[3, 30], phenylalanine ammonia-lyse[3, 31, 32], GSTs [3, 33-35], 14-3-3 like protein GF14-F/Drought induced protein 3/Peroxidase/Phosphoserine aminotransferase [29], ATP synthase [3, 6, 22, 25, 29], Cold shock domain protein [36], Drought-induced S-like ribonuclease [27], DUF26-like protein/Photosystem related proteins/Non-specific lipid-transfer protein [6, 29], Malate dehydrogenase [22, 37].

Based on physiological functions analysis of cold responsive proteins in previous studies, some proteins identified in the present work maybe have the function of resisting to cold stress. For example, 14-3-3 proteins regulate target proteins which involved in responses to biotic and abiotic stress through protein interactions [38-40], rice GF14c can target to plasma, thylakoid and vacuolar membranes and associated ATPase synthase complexes involved in stress responses [41-43]. 14-3-3-like protein GF14-F was shown up-regulated during cold stress in this study, and is probably function for cold resistance in the cold tolerance rice cultivar Kongyu131. Calcium-transporting ATPase genes differentially expressed under cold, salt and drought stresses involved in abiotic stress triggered signaling [44], the protein up-regulated after 72 h cold stress treatment maybe trigger stress signaling pathway. Chaperone protein ClpD1 is involved in heat and osmotic stress response, and the protein up-regulation is correlated with increased drought tolerance in rice [45], the protein up-regulated both at 48 h and 72 h cold stress treatments maybe play role in resisting to cold stress in this experiment. 6-phosphogluconate dehydrogenase activity increased in rice seedlings during various abiotic stresses treatments, maybe function as a regulator to control the efficiency of the pathway under abiotic stresses [46]. Cold shock domain proteins can be inducible expression under cold stress conditions in plants for cold acclimation, such as *Arabidopsis* and winter wheat [47-49]. Cold shock domain proteins shown to be not accumulate during the low temperature stress treatment in rice [36], however, the protein level decreased during cold stress treatment in this study (Table 1-3), maybe the protein has different regulation in different rice varieties which have different resistant levels to cold stress, these correlative data support the notion that the protein maybe involved in the cold acclimation response.

Interesting, in the 289 DEPs, only 11 proteins continue up-regulated and 11 proteins continue down-regulated from 24 h to 72 h cold stress treatment. These continued regulation proteins during cold stress maybe play important role for rice cold resistance. Cell division cycle protein 48 (CDC48) associated with leaf senescence and plant survival in rice [50], and a single base substitution on CDC48 can change the sensitivity of yeast to cold stress and cause cell death [51]. A homolog of AtCDC48, AtOM66 which located to the outer mitochondrial membrane in *Arabidopsis* plays a role in regulating cell death in response to biotic and abiotic stresses [52]. In present study, CDC48 protein up-regulation maybe play role in enhancing survival of Kongyu131 by determining the progression of cell death during cold stress treatment. In previous study, glycolate oxidase (GLO) maybe interact with catalase (CAT) to regulate H_2_O_2_ levels in rice under environmental stress or stimuli [53]; GLO5 up-regulation in this study, it may play role in cold resistance of rice. Although other DEPs identified in the work were not be reported have function for cold stress response, these proteins up- or down-regulated under cold stress are now able to be annotated as “cold-regulated proteins”. The physiology functions of these DEPs will be needed to fully characterize in further studies to enhance our understanding of cold stress responses in plants at the molecular level.

## Conclusion

In conclusion, this study is the first to adopt iTRAQ-based quantitative proteomics approach to identify cold response proteins in cold-tolerance *japonica* rice cultivar Kongyu131. A total of 289 cold responsive proteins were identified in time courses cold stress treatment, and the number of DEPs increased with the cold stress treatment time increased. Partial cold responsive proteins which have function in abiotic stresses were also identified in this study, such as 14-3-3 proteins, cold shock domain protein, CDC48 protein and GLO5. Some unknown proteins were first be identified in this study, specially continue up-regulated proteins (Q0JN91, Q6YU90, B9F813, A0A0P0Y5F2) and continue down-regulated proteins (A0A0N7KH91, Q0D5S1) from 24 h to 72 h cold stress treatment, maybe these proteins have important function for cold tolerance of rice variety Kongyu131. In this study, the DEPs were not be identified in previous studies of cold-treatment rice provides candidate proteins for further biological function study to better understand the cold-tolerance mechanism of rice responses to cold stress.

## Supporting information

**S1Table. Identified information of rice proteins by iTRAQ labeling**.

**S2 Table. Primer information using for Q-PCR**.

## Acknowledgments

This work was supported by the National Natural Science Foundation of China (grant no. 31960059, 31960401), Guangxi Natural Science Foundation (grant no. 2017GXNSFAA198266, 2018GXNSFAA050128), Guangxi Science and Technology Base and Special Talents (grant no. GuiKe AD18281069, GuiKe AD18050002) and Science-Technology Development Funding of Guangxi Academy of Agricultural Science (grant no. 31960059, 2021JM23, GuiNongKe2020YM124).

## Abbreviations

iTRAQ: Isobaric tags for relative and absolute quantification

DEPs: Differentially expressed proteins

GO: Gene Ontology

KEGG: Kyoto Encyclopedia of Genes and Genomes

Q-PCR: Quantitative real time PCR

FDR: False discovery rate

